# Eye movement markers of relational encoding predict scene memory precision

**DOI:** 10.64898/2026.09.30.754408

**Authors:** Eleanor Alderman, Eleanor Carre, Matteo Lisi, Szonya Durant, Carl J. Hodgetts

## Abstract

Recalling scenes and events depends on how information is relationally organised in memory, but it is unclear whether the organisation of eye movements during encoding, rather than simply how much is explored, shapes what is remembered. Using a scene-viewing paradigm and drawing-based free recall task to capture the content and structure of remembered scenes, we examined whether scene memory was better explained by ‘lower-order’ measures reflecting the magnitude and spatial extent of sampling, or ‘higher-order’ measures capturing gaze organisation and temporal dynamics. Relative to lower-order measures (number of fixations and scanpath length), memory precision was better predicted by repeated sampling of viewed regions (refixations) and gaze distributions (stationary gaze entropy) during encoding. Thus, the organisation of visual attention, rather than sampling quantity, contributed to scene representations that preserved relationships between elements. These findings highlight a role for spatial gaze dynamics in optimally encoding organised, relational scene representations for episodic memory.

---

Real-world scenes are thought to provide the ‘backdrop’ to our episodic memories – our memory for personal past experiences – providing the spatial and situational context to all events we encounter in our day-to-day lives (Armson et al., 2021; Robin, 2018; Robin & Olsen, 2019; Rubin et al., 2019). Scenes, by definition, consist of multiple objects organised hierarchically with respect to some background layout; therefore, their perception requires disparate but interconnected elements to be visually sampled and bound into a coherent percept (Castelhano & Henderson, 2008; Castelhano & Krzyś, 2020; Williams & Castelhano, 2019). As such, visual exploration of an environment can be thought of as the first step of encoding, where we initially process and integrate visual information into memory and form coherent internal representations of naturalistic spaces (Hollingworth, 2004; Ryan & Shen, 2020).

Scene perception typically relies on two forms of eye movement: fixations, the discrete stops our eyes make during which visual information can be processed (Castelhano & Rayner, 2009); and saccades, the frequent rapid shifts we make with our eyes between two fixations (Foulsham, 2015; Henderson & Hollingworth, 2003b; Land, 1999; Richardson et al., 2007). This allows us to direct our attention and focus with high visual acuity to relevant parts of the scene; facilitating the formation of detailed internal representations of real-world visual scenes (Henderson, 2003, 2017; Rayner, 2009; Yarbus, 1967). Such visual behaviours during scene perception may therefore provide insight into the process of constructing visual scene representations for memory.

Previous research has linked a greater number of fixations during encoding with enhanced subsequent memory across a range of naturalistic visual stimuli (Loftus, 1972; Henderson et al., 2005). This association has also been observed specifically during scene encoding, with individuals who make a greater number of fixations showing enhanced subsequent written free recall (Fehlmann et al., 2020) and recognition memory performance (Damiano & Walther, 2015, 2019). Likewise, a greater breadth of visual sampling across scenes has been associated with improved scene memory recognition success (Damiano & Walther, 2019; Mikhailova et al., 2021; Olejarczyk et al., 2014), and greater fixation density on objects within a scene corresponds to greater object memory performance (Celli et al., 2025). Consistent with this, restricting eye movements during encoding impairs subsequent memory for two-dimensional (2D) naturalistic scenes (Liu et al., 2020), suggesting that visual exploration is actively contributing to scene memory formation. Together, these findings suggest that measures reflecting the extent of visual sampling provide an important behavioural index of successful scene encoding.

Whilst measures such as the amount of eye movements during encoding are informative indicators of visual sampling, they focus on the isolated, discrete aspect of eye movements and do not capture the relationship between these fixations per se. In other words, such ‘lower-order’ visual metrics reflect the extent of visual exploration, but not *how* one’s gaze is organised over time or across space. Scenes are fundamentally structural, containing complex hierarchical relationships between larger global scene-defining features and smaller-scale objects (e.g., the computer is on top of the table and next to the keyboard in the corner of the office; Kaiser et al., 2020; Sun et al., 2025). However, despite the inherently relational structure of scenes, there is relatively little known about whether we adopt relational patterns of eye movements during scene processing and whether such patterns facilitate the construction of detailed memory representations that preserve the relationship between scene elements (e.g., comparing, updating, or binding information across locations; Eichenbaum, 2017; Rubin & Cohen, 2017).

Capturing the relationship between fixations, therefore, may be essential for understanding the dynamic nature of scene viewing and for furthering our understanding of precisely how different mechanisms of exploration facilitate the encoding and recall of scene environments. For example, gaze refixations – defined as fixations to a previously viewed region – are a common exploration pattern used during visual perception and scene visual search tasks (Kragel & Voss, 2022; Nikolaev et al., 2025b; Meghanathan et al., 2019; Rondina et al., 2016; Zelinksy et al., 2011; Zhang et al., 2022). Specifically, revisiting an area previously looked at is thought to be a key component of forming a comprehensive representation of the space by resampling areas of a scene, which could in turn lead to greater memory for the space globally (Kragel & Voss, 2022; Nikolaev et al., 2025a; Smith & Henderson, 2009; Sun et al., 2025). Also, as refixations are, by their definition, repeated sequential sampling of different areas of the scene, they are often considered key for processing relational structure within a visual scene (Kafkas & Montaldi, 2011). Refixations may also be a key component of visual working memory during exploration over time, whereby returning the gaze to regions already fixated within a scene can act as a rehearsal mechanism to reduce immediate decay and facilitate in-the-moment scene processing (Zelinsky et al., 2011). Notably, a greater number of refixations during the initial encoding of scene images can lead to better memory recognition performance (Holm & Mäntyla, 2007; Kragel et al., 2021), providing evidence that how we fixate between regions of a scene may facilitate our memory processes.

Additionally, measures of gaze entropy reflecting the complexity of fixation locations (stationary gaze entropy: dispersion across an image; gaze transition entropy: the degree of uncertainty from one fixation to the next) have been shown to differ depending on task complexity (e.g., driving, surgeons, aviation; Allsop et al., 2016; Di Stasi et al., 2016; Schieber & Gilland, 2008), suggesting they may reflect detailed in-the-moment processing and engagement with one’s environment. Therefore, summarising relationships among fixations in this way can be thought of as ‘higher-order’ metrics – providing a measure of how, and not simply to what extent, individuals distribute their attention when engaging with a task or new environment.

When recalling both episodic memories, and scenes, we are reconstructing a series of spatial- and/or temporal-relational elements that are bound together into a coherent event or percept. Therefore, relational information is not only key for scene perception, but critical for how scenes and events are recalled (Robin et al., 2016; Robin, 2018). Previous research predominantly relies on recognition-based memory tasks (e.g., Damiano & Walther, 2015, 2019; Olejarczyk et al., 2014; Mikhailova et al., 2021), in which encoding-related eye movements are related to a binary old/new judgement at test. As recognition can be supported by familiarity or gist-based signals alone, without access to the spatial-relational detail thought to underlie recollection (Castelhano & Krzyś, 2020; Cowell et al., 2019; Rubin & Cohen, 2017), this approach cannot inform whether higher-order relational eye movements predict the content and fidelity of scene memory – that is, how much spatial-relational detail is retained, and how accurately – as opposed to simply whether a scene feels familiar.

An alternative method for evaluating memory, particularly in accessing one’s internal representations, is through drawing (a ‘visual recall task’; Bainbridge et al., 2019). Scene drawing tasks have recently been implemented to assess individuals’ memory for spatial environments, including scene images (e.g., Bainbridge et al., 2019; Hall et al., 2022), and virtual environments (Cen et al., 2024). Notably, recollected drawings have been found to be representative of the layout of features and objects within the original scene images, and were not solely based on participants’ pre-existing schema of the scene categories presented (Bainbridge et al., 2019, 2021; Hall et al., 2022). Therefore, such scene drawing measures, when coupled with higher-order eye movement metrics, may allow us to examine how the organisation and temporal dynamics of scene visual exploration relate to what objects and structural features are successfully encoded, and how these features are recalled relative to one another – something that standard recognition or written recall tasks are unable to capture.

To examine whether the content and organisation of scene memory reflected visual exploration during encoding, we used a scene drawing recall task with detailed manual coding of object and spatial-relational features (see Methods and Fig. 1), capturing the accuracy and detail of recollected object locations, spatial relationships between key features and objects, and global scene structure (Bainbridge et al., 2019; Cen et al., 2024). We first assessed whether fixated object regions were more likely to be recalled than non-fixated regions, predicting that objects fixated during encoding would be more likely to be successfully remembered at recall. Based on previous evidence linking gaze behaviour during encoding with subsequent memory, we further predicted that lower-order eye movement measures reflecting the amount of visual exploration (fixation count, scanpath length) would positively predict memory precision scores during the scene drawing recall task. We also examined whether relational patterns of eye movements related to subsequent scene memory precision, predicting that higher-order measures (refixations, stationary gaze entropy, gaze transition entropy) would positively predict memory precision scores, reflecting how the distribution of attention during 2D scene viewing relates to subsequent recollection. Critically, given that higher-order measures capture the relational structure thought to underlie recollection (see above), we predicted that higher-order measures of gaze organisation would significantly predict memory precision.

**Fig. 1.**
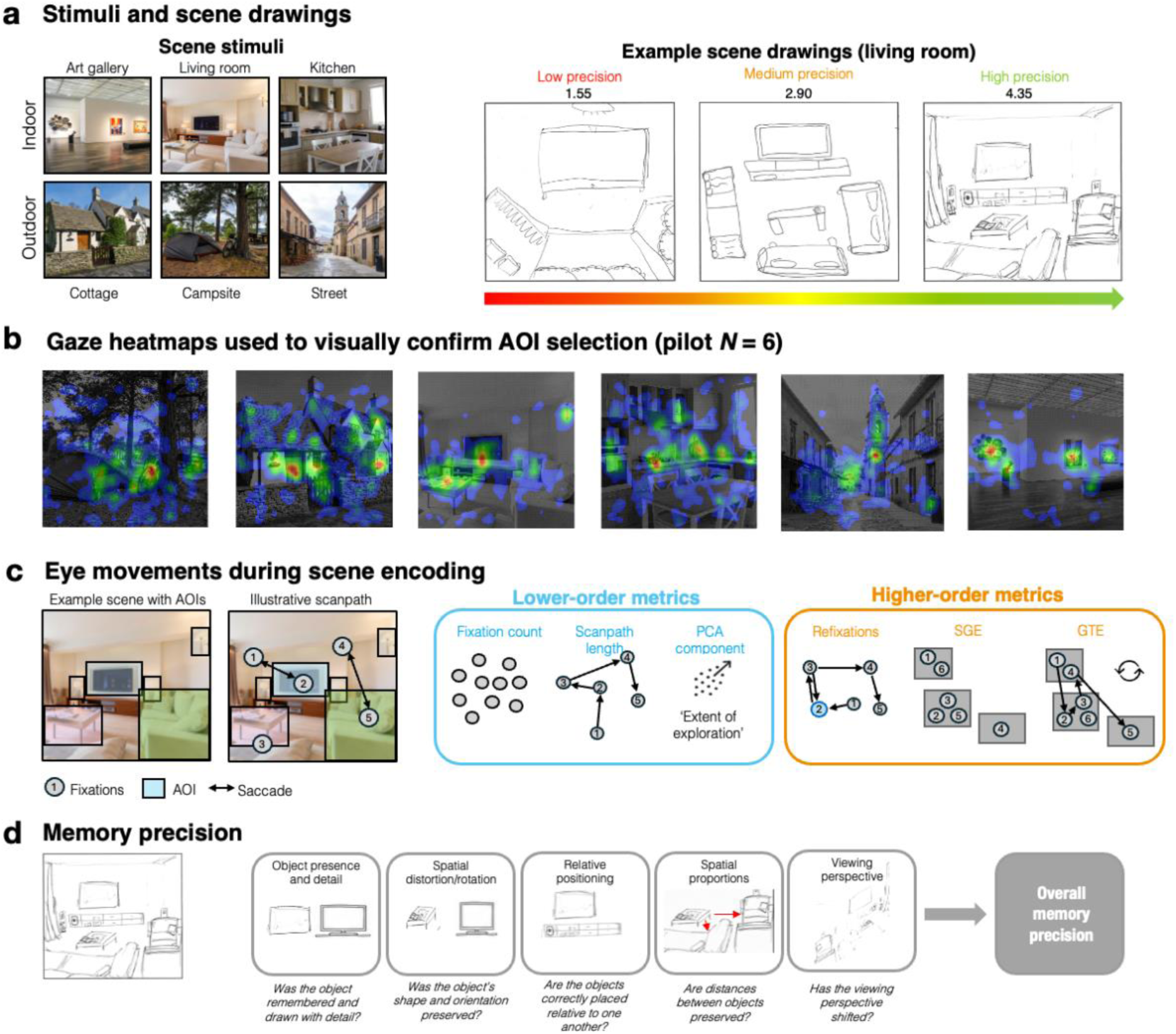
An overview of the experimental approach, eye movement metrics, and memory coding scheme. (A) All six scene images (3 indoor; 3 outdoor) presented to participants at encoding, including examples of low, medium, and high precision drawings of the living room from three participants. The scores given to each drawing is shown. (B) Gaze heatmaps from pilot participants used to verify defined object AOIs. Red to blue gradient represents the total number of gaze samples at each location scaled for each image, with red indicating a higher number of gaze samples. (C) Eye movement metrics in the present study, including schematic of scanpath across a scene image. Depictions of all defined gaze metrics are provided, categorised based on lower- and higher-order behaviours. (D) Examples of the coding scene, including manipulated exemplar drawing from a participant to represent different scored metrics from the coding scheme. All coding items were manually scored by two raters. See ‘Scoring of memory drawings’ for reported ICC values.

We additionally explored whether subjective aspects of recall were associated with drawing-based memory performance. First, we assessed whether retrospective confidence in recall accuracy corresponded with subsequent memory precision. Self-reported confidence in believed recognition performance accuracy has been shown to relate to actual recognition performance accuracy (Greene et al., 2024), suggesting that individuals can demonstrate insight into the quality of their own memory representations. An association between confidence ratings and memory precision would therefore suggest that our task captures meaningful variation in memory fidelity. We also explored whether subjective vividness and immersion during the scene drawing task related to subsequent memory. A greater sense of presence during episodic memory recall (Smith & Mulligan, 2021) and the recollection of encoded movie stimuli (Makowski et al., 2017) has been linked to enhanced recall abilities, suggesting that the subjective experience of engaging with a scene may influence the formation and recollection of memory representations. These exploratory analyses provide further insight into whether the perceived richness and immersion of encoded scene representations relates to memory precision.

## Results

To explore the role of visual exploration behaviours in scene memory, we sequentially presented participants with six real-world scene images (three indoor; three outdoor) for 20 seconds each, while tracking their eye movements (see Methods for further detail). We then assessed the precision of their scene memory using a drawing paradigm, where participants freely recalled as many of the previously viewed scenes as possible, with unconstrained time to do so (see Fig. 1A for example drawings). We then derived several measures of visual exploration from the eye tracking data: ‘lower-order’ eye movement metrics reflecting the extent of visual exploration (fixation count, scanpath length), and ‘higher-order’ measures (refixations, stationary gaze entropy, gaze transition entropy) reflecting the spatial organisation and temporal dynamics of gaze (shown schematically in Fig. 1C). To measure memory precision, we developed a coding scheme (adapted from Cen et al., 2024) to assess object and spatial-relational detail within participants’ scene drawings. For our main analyses, we then ran three linear mixed effects models (LMMs; see Methods for detailed model specifications). First, we tested whether attention to individual objects (looked at or not) was reflected in their later recall (present or not). Second, we explored how lower-order metrics across scene images during encoding may relate to overall scene precision. Third, we explored whether higher-order, relational patterns of eye movements (e.g., the predictability and distribution of gaze across a scene image) related to overall scene memory precision. We predicted that greater visual exploration overall, reflected by both lower-order and higher-order metrics will be predictive of overall scene memory precision scores.

### The distribution of eye movement and memory metrics

We first characterised the distribution of our main eye movement and memory metrics, as well as between-scene differences. On average, participants spent approximately 12min 36s minutes (SD = 4min 59s), completing the scene drawing task. Additionally, scores from participants’ copy drawings of the campsite were calculated using the same coding procedure (see Table 1 for descriptive statistics). One participant obtained a notably low copy drawing score of 2.85, which was visually inspected and indicated sufficient engagement with the task, and was included in final analyses. Fig. 2 displays scene-level variability across memory measures and distribution of overall memory precision scores. Although mean values varied across scene images, distributions showed substantial overlap and comparable levels of within-scene variability. To account for potential stimulus- and participant-level differences, subsequent analyses incorporated scene and participant as random effects within LMMs, and all participants retained in the analysis contributed observations from at least four of the six scene images. Full distribution across our eye movement metrics are shown in Supplementary Fig. S1.

**Fig. 2.**
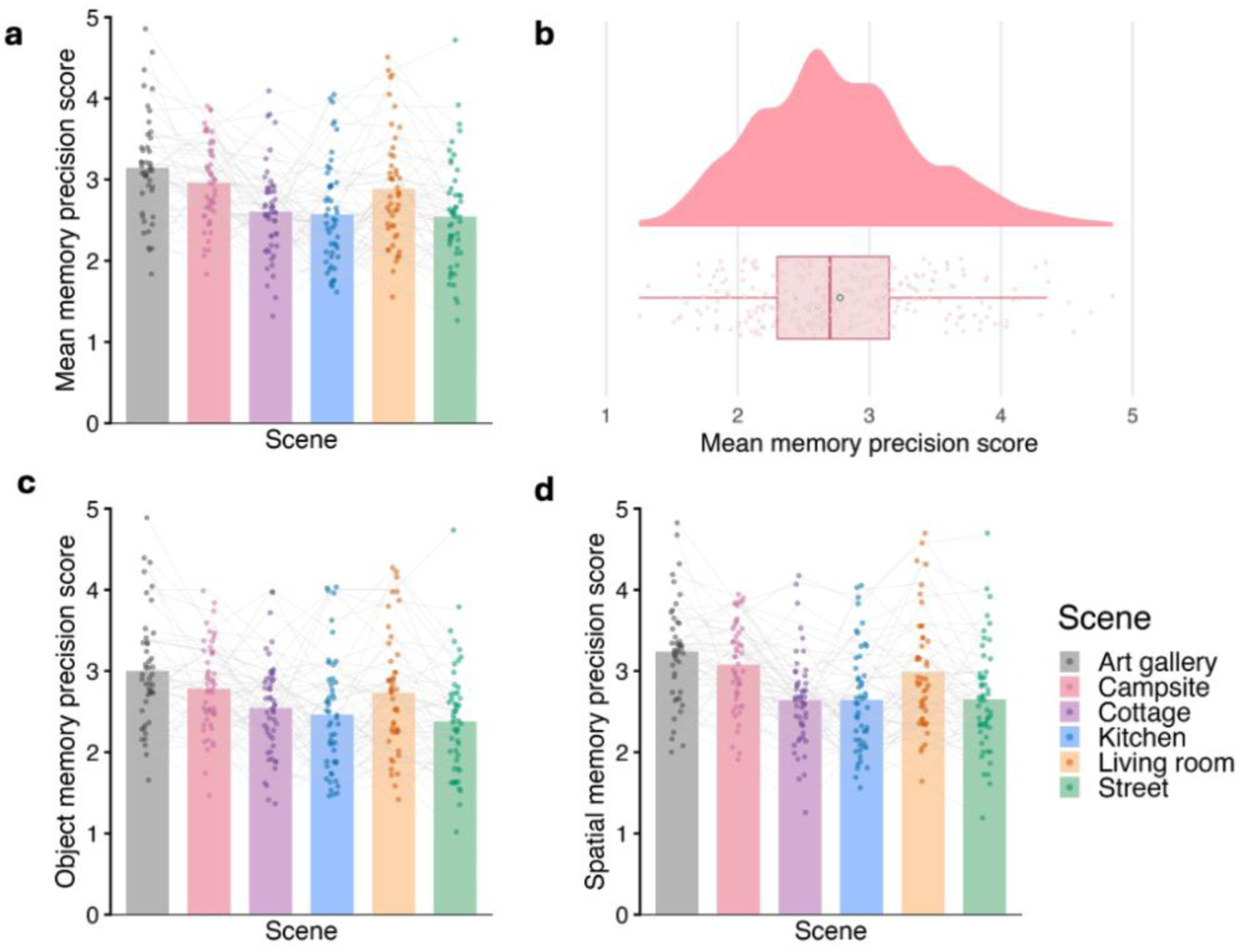
Descriptives for overall object, and spatial memory scores, broken down at scene-level and overall memory score distribution. Jittered points represent individual observations, with lines connecting dots between individual participants.

**Table 1.**
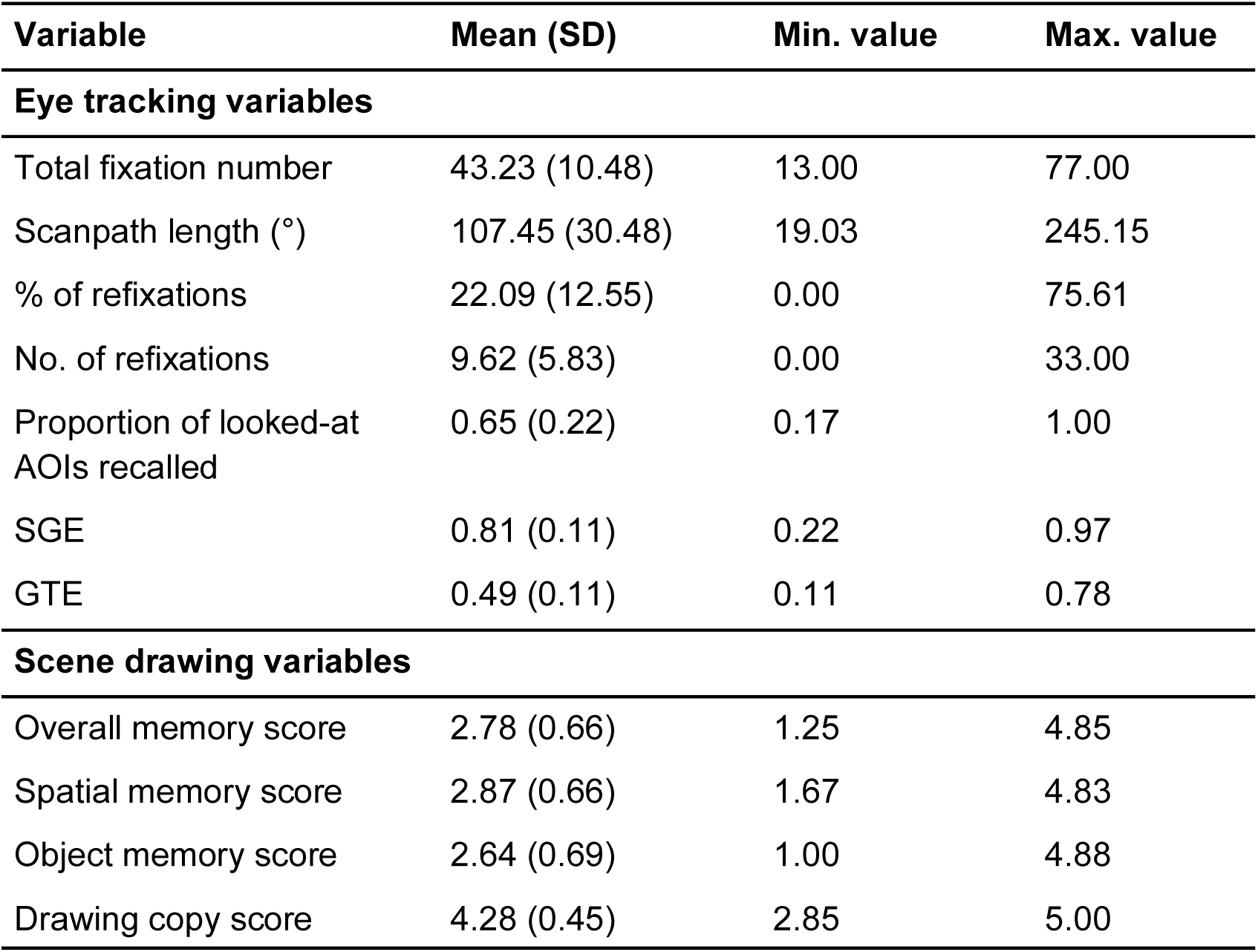
Descriptive Statistics (Mean, Standard Deviation, and Range) for Eye Tracking and Memory Variables of Interest.

#### Fixating on scene AOIs during encoding predicts their later recall in scene drawing task

To test whether fixating upon a defined scene region of interest during encoding increases the probability of later recalling it, we fitted a generalised linear mixed effects model (GLMM). The model revealed a significant effect of fixation on recall, whereby AOIs fixated on during encoding were more likely to be recalled than AOIs that were not (*β* = 1.52, *SE* = 0.18, *z* = 8.57, *p* < .001, OR = 4.59 95% CI [3.27, 6.63]). Additionally, this corresponded to a predicted recall probability of ∼66% for fixated AOIs, compared to ∼29% for non-fixated AOIs. This finding supports the initial hypothesis that fixating on a particular scene region at encoding supports subsequent memory.

#### The extent of visual exploration does not predict scene memory precision

We next asked whether the amount of visual exploration – i.e., how broadly participants scanned a scene – predicted the fidelity of what is later recalled during our scene drawing task. Addressing this, we fitted an LMM using a PCA-derived component capturing extent of visual exploration (fixation count and scanpath length; see Methods) as a fixed effect to assess its predictive effect on memory precision, going beyond the analysis above Model 1 to capture both spatial and object detail captured in recollected scene drawings. We observed that the extent of visual exploration, as captured by our PCA composite score, was not a significant predictor of overall scene memory precision (*β* = -0.001, *SE* = 0.029, 95% CI [-0.063, 0.055], *t*(284) = -0.036, *p* = 0.971; Fig. 3A). This suggests, therefore, that the amount of precise object and spatial-relational detail recalled is not driven simply by the degree and extent of visual sampling.

**Fig. 3.**
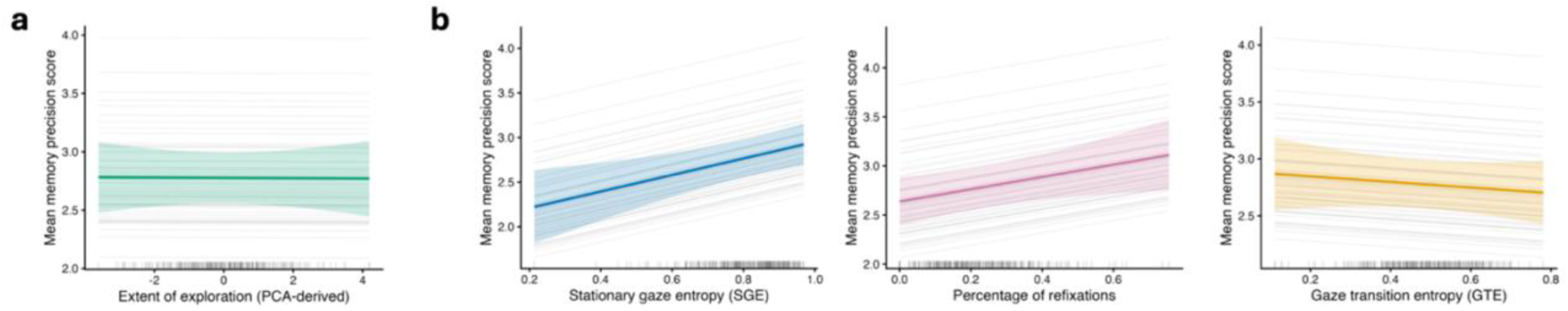
Associations between lower- and higher-order predictors and mean memory score. (A) Linear mixed effects model estimates showing the associations between mean memory score and extent of exploration (‘lower-order’). (B) Linear mixed effects model showing the associations between gaze transition entropy, stationary gaze entropy, and percentage of refixations (‘higher-order’), respectively. Grey lines represent individual participant relationships and coloured lines represent group-level model estimates with shaded areas showing 95% confidence intervals. Vertical grey lines show the distribution of scores across participants. Grey marks along the x-axis indicate the distribution of observed predictor values.

#### Relational gaze patterns predict memory precision

Testing our key hypothesis, we next examined how the distribution of gaze (SGE, GTE) and relational patterns of eye movements across a scene (refixations) related to overall memory precision. An LMM including GTE, SGE, and percentage of refixations as predictors revealed that SGE (*β* = 0.95, *SE* = 0.30, 95% CI [0.350, 1.434, *t*(262) = 3.23, *p* = 0.001; Fig. 3B) and percentage of refixations (*β* = 0.62, *SE* = 0.28, 95% CI [0.062, 1.174], *t*(274) = 2.23, *p* = 0.027; Fig. 3C) both positively predicted memory precision scores, whilst GTE did not (*β* = -0.28, *SE* = 0.32, 95% CI [-0.917, 0.300], *t*(259) = -0.88, *p* = 0.378; see Fig. 3D). Participants who distributed their fixations more evenly across scene regions and who returned to previously-viewed locations more frequently (which is proposed to reflect more hierarchical relational processing) subsequently reproduced scenes with greater object and spatial fidelity.

This model accounted for 54.1% of overall variance in memory precision scores, with 3.3% of variance being explained by the fixed effects predictors (conditional *R^2^*= 0.541, marginal *R^2^* = 0.033). Whilst much of the variance was due to differences between participants and scenes as suggested by the random effects of participant (variance = 0.174, SD = 0.417) and scene (variance = 0.054, SD = 0.232), the positive association here suggests that a greater percentage of refixations, and increasing SGE was associated with enhanced memory precision as demonstrated by scene drawing scores. Together, these findings suggest that relational and redistributive gaze behaviours – rather than raw exploratory volume or sampling rate – are key eye movement signatures of successful and spatially coherent encoding.

To compare relative fit of the lower-order and higher-order eye LMMs, both models were first refit using maximum likelihood (ML) estimation and were compared using Akaike’s Information Criterion (AIC), with lower values indicating better relative model fit. The higher-order eye movement model provided a better fit to the data than the lower-order model (higher-order AIC = 491.9, lower-order AIC = 502.9), indicating greater support for the higher-order model as an explanation of overall subsequent scene memory precision.

### Exploratory analyses

#### Subjective memory experience predicts scene reconstruction precision

To characterise the phenomenology of recall whilst drawing the scene images, participants were also asked to rate their subjective confidence that their drawing accurately represented the original scene, the vividness of the mental image during recall, and their sense of spatial immersion during recall (i.e., to what degree they visualised the scene image from a 2D versus 3D perspective; see Table 2 for subjective rating descriptives and Supplementary Fig. S2 for distribution scores of all exploratory variables). To understand how these subjective ratings relate to scene memory precision, we ran an exploratory LMM with confidence, vividness, and immersion as fixed effect predictors. Participant and scene were entered as random effects in the model.

**Table 2.** Descriptive Statistics (Mean, Standard Deviation, and Range) for Subjective Judgement Ratings.

| Variable | Mean (SD) | Min. value | Max. value |
| --- | --- | --- | --- |
| Confidence | 2.82 (0.96) | 1 | 5 |
| Vividness | 53.60 (22.29) | 0 | 100 |
| Immersion | 45.60 (25.56) | 0 | 95 |

Overall, both vividness during recall (on a scale of 1-100; *β* = 0.0049, *SE* = 0.0018, *t*(271) = 2.67, *p* = 0.008, 95% bootstrap CI [0.002, 0.008]; see Supplementary Fig. S3A) and confidence in one’s recalled drawing (on a scale of 1-5; *β* = 0.2060, SE = 0.0390, *t*(278) = 5.28, *p* < 0.001, 95% bootstrap CI [0.128, 0.292]; see Supplementary Fig. S3B) were significant predictors of scene memory precision. Specifically, a one-point increase in vividness ratings was associated with a 0.005 increase in scene memory precision scores, whereas a one-point increase in confidence ratings led to a 0.21 increase in scene memory precision scores. Sense of immersion was not found to be a significant predictor in the model (*β* = 0.0013, *SE* = 0.0016, *t*(288) = 0.845, *p* = 0.399, 95% bootstrap CI [-0.003, 0.005]; see Supplementary Fig. S3C). Overall, this model explained 63% of the explained variance in the model, with 20.4% of the variance being explained by subjective experience measures. These exploratory analyses suggest that how vividly and confidently a participant mentally re-experienced a scene was meaningfully tied to how accurately they could reconstruct it.

#### No evidence of boundary extension in scene recall

Scene drawings were also coded for boundary extension, the tendency to recall scene content beyond the original image boundary, reflecting a well-documented bias toward extended spatial representations in memory (Bainbridge & Baker, 2020). Boundary extension was negligible across participants (*M* = -0.02, *SD* = 0.26; see Supplementary Fig. S3D), with no reliable deviation from zero (one sample (one-sample *t*(296) = −1.45, *p* = 0.926) and was not a significant predictor of memory precision (*β* = −0.052, *SE* = 0.126, *t*(275) = −0.413, *p* = 0.680; see Supplementary Fig. S3E. This suggests that participants tended to draw accurately within the implied scene boundary, with little systematic tendency to extend beyond (or indeed compress) the boundary of the encoded image.

#### False memories are associated with lower scene reconstruction precision

Drawings were additionally coded for false memories, capturing recalled objects or features absent from the reference image. Although overall false memory rates were low (*M* = 1.18, *SD* = 0.29), it was a significant negative predictor of memory precision (*β* = −0.427, *SE* = 0.110, *t*(272) = −3.880, *p* < 0.001; see Supplementary Fig. S3F), accounting for 3.7% of fixed-effects variance in an LMM that explained 53.6% of total variance overall. Therefore, the tendency to reproduce false details at recall was associated with poorer overall recollection of object and spatial features of the reference scene.

## Discussion

While prior work has demonstrated that visual exploration during scene perception can support subsequent memory success, little is known about how the relational organisation of visual exploration relates to the precision of scene memory. By combining eye tracking with a free recall scene drawing task, we quantified both lower- and higher-order eye movements reflecting extent of exploration and spatial-relational exploration, respectively, to examine how different gaze behaviours support the recall of object and spatial details. First, we found that participants were more likely to recall object AOIs if they were fixated on during encoding, consistent with prior research suggesting that fixating on objects can lead to stronger internal representations of such targets (Castelhano & Henderson, 2005; Henderson & Hollingworth, 2003a; Hollingworth, 2004). Second, we observed that lower-order measures, combining number of fixations and scanpath length into one PCA-derived variable, were not found to significantly predict memory precision scores, suggesting that the extent of visual exploration at encoding did not influence the content and organisation of scene recall as assessed via scene drawings.

Aligned with our predictions, a model containing GTE, SGE, and percentage of refixations significantly predicted memory precision. Whilst GTE itself did not predict memory precision scores, higher SGE and a greater percentage of refixations – indexing a more relational and distributed allocation of gaze – were both associated with enhanced memory precision scores. These findings suggest that the redistribution of fixations across scene regions, and not the specific order and predictability in which gaze moves between locations (as reflected by GTE), may contribute to the formation of more precise scene representations. Model comparisons revealed that this higher-order model better explained the data than our lower-order model, suggesting the relational measures of eye movements better captured the variability in scene memory precision than measures reflecting the extent of visual exploration.

Whilst previous research has consistently reported positive associations between the amount of visual exploration and scene memory (e.g., number of fixations; Damiano & Walther, 2015, 2019; Fehlmann et al., 2020; Olejarczyk et al., 2014), this relationship was not observed in the present study. To our knowledge, this is the first study to examine how encoding-related eye movement metrics predict performance in a full visual scene recall task, where recall required participants to reconstruct specific object details and holistic spatial relationships with accuracy and precision (e.g., Bainbridge et al., 2019). Compared to recognition paradigms, this task places greater demands on the content and fidelity of the encoded representation, as successful memory cannot be supported by familiarity judgements alone (Yonelinas, 2002; Yonelinas et al., 2010). This difference in task demands may explain this disparity in findings, as sampling alone, without requiring the same degree of gaze organisation across time and space, may not be sufficient for precise memory reconstruction.

Consistent with this account, our findings align with evidence that relational gaze behaviours (e.g., refixations) facilitate memory performance (Kragel et al., 2021; Kragel & Voss, 2022). Rather than reflecting simply how much information is sampled, higher-order measures may capture how visual attention is organised across a scene, supporting the binding of features into coherent representations. This may be particularly beneficial for precise recall, whereas fixation count may suffice for less detailed retrieval (e.g., gist-based recognition or verbal recall). Higher-order gaze behaviours, including refixations, may also reflect online cognitive processing during scene exploration. Refixations may help to offset interference from subsequent fixations and maintain information in visual working memory (Castelhano & Henderson, 2005; Meghanathan et al., 2019; Zelinsky et al., 2011), while individuals who spontaneously make more refixations during encoding tend to make fewer spatial memory errors (Kragel et al., 2021). Refixations may therefore reflect the resampling of insufficiently sampled scene regions, allowing integration of spatial relationships between scene components, strengthening relational representations and allowing greater memory precision during recall (Kafkas & Montaldi, 2011). This result posits refixations as a potential marker for relational processing.

Moreover, SGE provides an indication of the spatial organisation of gaze across the scene, and has also been associated with task complexity and engagement, consistent with theories suggesting that successful scene recall depends on complex hierarchical representations of a scene (Maguire & Mullaly, 2013; Robin et al., 2016; Robin & Olsen, 2019), including encoding spatial relationships between key features. Thus, refixations and SGE may reflect aspects of gaze organisation that support the construction of spatial representations, preserving information about object relationships and broader scene structure.

This interpretation aligns with relational memory theories, which propose that successful memory depends on binding individual elements into structured representations, and is hypothesised to depend on the hippocampal system (Eichenbaum, 2017; Rubin & Cohen, 2017). Further, given the proposed role of the hippocampus in scene-based cognition (Lee et al., 2012; Murray et al., 2017; Zeidman & Maguire, 2016), these relational gaze behaviours may relate to hippocampally-supported scene representations during encoding (Kragel et al., 2021; Kragel & Voss, 2022; Liu et al., 2018). Together, these findings suggest that the organisation of visual exploration may support the formation of precise scene representations by integrating spatial information within memory.

This work may also have applications for understanding memory in ageing and neurodegenerative disease. Prior work has shown that ageing alters eye movement patterns during encoding, even when recognition memory performance remains comparable to younger adults (Mazloum-Farzaghi et al., 2023), with older adults typically making more fixations during encoding (Firestone et al., 2007; Heisz & Ryan, 2011; Ryan et al., 2007; Wynn et al., 2021). It has been proposed that increased sampling in older adults reflects up-regulation of hippocampal relational binding processes (Liu et al., 2018). Given that scene drawing and higher-order eye movement metrics may be particularly sensitive to relational encoding processes, our results may provide a basis for more detailed studies of perception and memory in ageing and neurodegenerative disorders.

As exploratory analyses, we examined whether retrospective subjective measures were associated with memory precision. Although self-reported sense of immersion during recall was not related to subsequent scene memory, greater vividness of the remembered scene was associated with higher memory precision. This aligns with previous research demonstrating that trial-by-trial stimulus-specific vividness of mental imagery for scenes is associated with subsequent memory accuracy (Duckett & Simons, 2026), suggesting that the subjective richness of recalled scene information may reflect the strength of the encoded memory representation. Greater confidence in recall accuracy was also associated with higher memory precision scores, consistent with evidence that individuals can show insight into the accuracy of their own memory performance across working- and long-term memory tasks (Greene et al., 2024). Together, these findings suggest that self-reported phenomenological experiences, including vividness and confidence, correspond with variation in scene memory precision.

Additionally, we explored whether memory distortions or errors within participants’ drawings were related to overall memory precision. False memory – the inclusion of incorrect objects or scene details – was associated with lower memory precision scores, suggesting that drawings containing more memory errors also tended to preserve fewer object and spatial-relational details. Moreover, despite previous reports that boundary extension is a common distortion in scene memory, including in drawing tasks (Bainbridge & Baker, 2020; Bainbridge et al., 2019), we observed little evidence of participants extending beyond scene boundaries, and boundary extension was not associated with subsequent memory precision. Previous research has suggested that boundary extension may depend on the type of scene initially presented, whereby wider-view scenes, such as those used in the current study, experience this fallacy to a lesser extent than close-up images (Hubbard, 2025). However, future work is needed to determine whether this reflects characteristics of the drawing paradigm or the encoded naturalistic scenes themselves.

Whilst the present study combines eye movements with a visual scene drawing recall task, providing a more granular assessment of memory that is sensitive to both object detail and relational structure, there are considerations for future research. Our research, and much prior research, relies predominantly on the perception and encoding of 2D scene images (Damiano & Walther, 2019; Fehlmann et al., 2020; Olejarczyk et al., 2014). Sense of immersion at recall was not a significant predictor of memory precision, which may suggest the task did not initiate recollection of the scene as one would from a real-world memory, in which scenes are re-experienced as an integrated whole. Real-world scene exploration involves head movements and locomotion, resulting in perspective shifts that 2D viewing cannot capture (Bischof et al., 2023), but are likely to influence scene memory. Future research using fully immersive three-dimensional environments will allow us to understand how lower- and higher-order gaze behaviours interact with exploratory processes to shape memory for real-world complex scenes.

Moreover, the present findings were derived from a relatively constrained set of naturalistic scenes, with six example scene images. This was implemented (based on piloting) to reduce overall memory demand, ensuring that most participants could recall some aspect of most scenes, allowing meaningful scoring of recollected scene drawings. Although scene image was modelled as a random effect, the extent to which gaze patterns reflecting relational encoding generalise across different environments remains an important question. Scenes vary substantially in their complexity, spatial structure, semantic content, and relationships between elements, all of which are likely to influence scene viewing behaviour and subsequent memory (Bylinskii et al., 2015; Kyle-Davidson et al., 2025). Future research should examine whether the association between gaze organisation and scene memory precision extends across a broader range of environments and visual stimulus types to understand whether these relationships are maintained across stimuli differing in their structural properties and processing demands.

Together, the findings from the current study demonstrate that memory for visual environments is shaped not only by the extent of visual exploration or where we fixate, but by how gaze is relationally structured across space during initial encoding. Consistent with this notion, fixation count did not predict memory precision, suggesting that measures of visual exploration do not fully capture the processes required for tasks demanding precise reconstruction of object and spatial information. Future research should examine how relational gaze behaviours support scene memory across more dynamic and immersive environments to determine how visual exploration supports structured scene representations. Overall, combining eye movement measures with scene drawing may provide an important framework for understanding how day-to-day environments are encoded and reconstructed, how these processes may change across the lifespan, and how they support the recollection of episodic events situated within those environments.

## Methods

### Participants

All participants were students or staff from Royal Holloway, University of London, were fluent English speakers, and had reported normal or corrected-to-normal vision, and no current neurological disorders. Sixty-five participants aged +18 years were recruited for this study. Data from nine participants were excluded due to having insufficient memory data (omitted more than two scene images during the recall task), failure of eye tracking calibration at the time of testing, or losing more than 25% of eye tracker samples, resulting in a final sample of 56 participants (mean age = 21.38, SD = 4.15, 45 female, 9 male, one non-binary, and one gender fluid). Participants were invited to take part in a research study that aims to understand how we remember everyday scenes and environments and were reimbursed with a £5 voucher, or undergraduate course credits upon completing their participation. Ethical approval was granted by the Research Ethics Committee of Royal Holloway, University of London (ethics approval code: 4066-2024-01-08-18-10-ZHJT011), all participants provided informed consent prior to participation.

### Materials

#### Scene Images

For the encoding task participants were presented with a randomised sequence of the same six two-dimensional (2D) images of naturalistic real-world scenes on a computer screen using Psychopy (version 2022.2.4; Peirce et al., 2019). Based on initial pilot testing, six images were selected to provide an appropriate level of memory demand, allowing meaningful variability in recall performance while minimising ceiling and floor effects. Three indoor scene images (living room, art gallery, kitchen) and three outdoor scene images (cottage, street, campsite) were selected (Fig. 1A). All scene images were selected for being neutral scenes (i.e., not associated with an obvious emotional context), and representative of everyday spatial environments. Additional inclusion criteria included: images not containing any famous landmarks/locations, text, or people; a space where one could feasibly imagine an event taking place, more than three visually distinct items (e.g., objects), and not being focused around one single object. Images were obtained taken from Adobe Stock (https://stock.adobe.com/uk/) under license. They were reproduced solely for the use of stimulus presentation during the encoding task of the study, and when used as reference images by the coders during scoring of recalled scene drawings. Images were cropped so that their width and height were equal in size (500 x 500 pixels). See Fig. 1A for scene images with corresponding example participant drawings, and supplementary Fig. S4 for low, medium, and high precision scene drawings for each category. Areas of interest (AOI) used for memory coding and data analysis for each scene were manually defined based on salient objects present within the scene image; the number of AOIs ranged between five and eight across scene images. All AOIs were verified to contain most of the object-focused fixations by visually inspecting initial eye tracking data from six pilot participants (see Fig. 1B for gaze heatmaps from pilot data used to visually confirm AOIs). Where AOIs overlapped (due to key object areas being close together), fixations landing within overlapping regions were counted as fixations to both AOIs when determining whether an AOI had been fixated and when calculating SGE, and were excluded from GTE calculations (see ‘Eye tracking’ below).

### Eye tracking

Participants’ eye movements were tracked using the Tobii TX300 eye tracker with a built-in 23” monitor (screen resolution 1920 x 1080). Participants’ gaze was sampled at 300 Hz. Before the encoding task began, the eye tracker was calibrated using a 5-point calibration, followed by a validation stimulus in PsychoPy (version 2022.2.4; Peirce et al., 2019). Participants were excluded based on their eye tracking data if their valid gaze samples for the duration of the scene viewing task, across all scene images, fell below the threshold (< 75%). Individual image trials falling below this threshold were also excluded. The monitor display subtended 42.9° horizontally at a viewing distance of 65cm, corresponding to 44.8 pixels/degree (0.022° of visual angle per pixel).

All eye tracking data were exported from PsychoPy and then processed using MATLAB R2024a (The MathWorks, Natick, MA, USA). Fixations were first identified using the I-VT (Velocity-Threshold Identification) filter, with a velocity threshold of 8 pixels per sample (2400 pixels/s), corresponding to approximately 53.6°/s (degrees of visual angle per second) based on the display size and viewing distance. Eye movements below this threshold were considered fixations. A minimum duration threshold of 60 ms was applied and Gaussian smoothing was applied to reduce noise in the raw gaze data (sigma = 6.7 ms). Missing samples were interpolated.

#### ‘Lower-order’ visual metrics

Lower-order eye movement metrics quantified the amount/depth of visual exploration during scene encoding. Total fixation count was defined as the total number of fixations made across the entire scene image during the encoding phase. Scanpath length was calculated as the sum of Euclidean distances between consecutive fixations. The number of AOIs fixated represented the number of predefined object regions of interest that received at least one fixation, with an AOI considered fixated if one or more fixation coordinates fell within its boundaries. Schematic illustrations of these metrics are shown in Fig. 1C.

#### ‘Higher-order’ visual metrics

Higher-order metrics capture the relational properties of eye movements, specifically, about the way visual information is sampled and integrated. Refixations were defined as fixations within 10 pixels (0.22° of visual angle) of any earlier fixation, indicating a return to a previously fixated region, but located more than 10 pixels (0.22° of visual angle) away from the immediately preceding fixation. This measure captured individuals’ tendency to revisit previously fixated regions during scene viewing. *Stationary gaze entropy (SGE)* and *gaze transition entropy (GTE)* were calculated to quantify the distribution and complexity of fixation locations and transitions across and between scene image AOIs. SGE measures the dispersion of fixation locations across AOIs, whereby higher SGE values indicate a more evenly distributed viewing pattern across the scene, whereas lower SGE values indicate a more concentrated viewing pattern (Ayala et al., 2023; Shiferaw et al., 2019). GTE captures the complexity and predictability of gaze transitions between AOIs, based on the extent to which subsequent fixation locations can be predicted from the location of the prior and subsequent fixation behaviours. Higher GTE values indicate a more complex and less predictable pattern of transitions between AOIs, whereas lower GTE values indicate a more consistent and predictable sequence of gaze shifts. SGE and GTE were calculated based on the supplementary materials provided by Shiferaw et al. (2019). Schematic illustrations of these metrics are likewise shown in Fig. 1C, and Matlab code for analysing eye tracking data is available on OSF: https://osf.io/c2xr9/overview?view_only=8e5af99b239d4e869088289f74beaa6c.

### Procedure

Upon arrival at the lab, participants were initially informed of the research purpose, and that the expected duration of this study would be 30-45 minutes. After providing informed consent, participants completed the encoding phase of the study. Participants were instructed to observe the images they would be presented with, and were additionally informed that their memory of these scenes will be tested, but the nature of this memory task was not disclosed. Once the eye tracker was calibrated and validated, the encoding phase of the study would begin. For the encoding phase, all six scene images in random order were shown to participants whilst their eye movements were tracked. Each image was presented in the centre of the display for 20 s, followed by a 1500 ms interstimulus interval, which consisted of a central fixation cross to re-centre the gaze in-between each image. Following this, participants then completed the recall phase of the study, where the instructions provided were to draw each image in ‘as much detail and as accurately as possible’. It was explained to participants that they should try to draw something for each image, even if they were not confident in their memory.

Once participants had completed their drawings, they then completed the subjective ratings measures for each scene image in turn. Here, participants were briefly presented each image in turn (the order of presentation was again randomised across participants) and were asked about their subjective experience of each image, being asked about their sense of immersion within the scene image, their confidence in the drawing they produced, and the vividness of the image in their mind during recall. Once all images had been presented, participants moved onto the copying phase of the experiment, whereby participants were instructed to draw the image of the campsite whilst the image remained on-screen. This served as a potential control measure for drawing ability and effort in the task paradigm. It was reinforced that this was not a test of memory or drawing ability, and they should focus on the objects and general structure of the scene and did not have to focus on including fine details in their drawings. Finally, participants were asked two final questions about their artistic abilities before having the opportunity to provide any comments about their experience of the drawing task. Participants were then fully debriefed regarding the nature and aims of the study and were then told that their participation in the study was complete.

### Memory coding and scoring

Scene drawings were coded using a scene memory coding scheme adapted from recent work on curiosity states and memory (Cen et al., 2024), which draws upon established approaches from human neuropsychology. This coding scheme assesses scene memory across both object-level (the accuracy of individual elements) and spatial-relational dimensions (the accuracy of overall scene structure) relative to the reference image (object metrics: *object presence and detail*, and *spatial distortion and rotation of objects*; spatial metrics: *relative positioning*, *spatial proportions*, and *viewing perspective*). Two exploratory items were also included to capture presence of *boundary extension* and *false memory* (all coding dimensions are outlined below).

Raters were also presented with a list of objects within each scene (corresponding to defined AOIs) and were required to select which of the objects were successfully recalled in the drawing – i.e., was there clear evidence that this object had been remembered, regardless of the accuracy in spatial location of its recall. All coding items were scored on a 9-point sliding scale from 1-5 (with 0.5 increments), where a higher score represents participants having better reflected that object or spatial element in their recollected drawing. Exceptions to this were the exploratory measures, whereby boundary extension was rated on a scale of -2 to 2 (to reflect both boundary contraction and boundary extension; -2 = ‘boundary contraction’ and +2 = ‘boundary extension’), and false memory, whereby 1 = ‘no false memory’, and 5 = ‘evidence of false memory’.

#### Object presence and detail

This criterion required the rater to score the number of objects within the scene drawing that were correctly recalled, and whether sufficient detail was captured in these items. Primarily, it aimed to assess whether there was evidence that the objects present within the original scene were recollected with accuracy in the scene drawing, regardless of accuracy of any other relevant spatial metric (e.g., positioning).

Here, a lower score reflected that none or very few of the original objects were included or objects lacked recognisable details, and a higher score indicated that all original objects were included with substantial detail.

#### Spatial distortion and rotation of objects

This item captured how well the dimensions, form, and orientation of individual objects and features were maintained and represented within the drawing. This was based on the objects that were correctly recollected, and drawings were not penalised if there were fewer correct objects recalled. If there was significant distortion or misorientation of objects or structural features within the scene, drawings were scored low on this item. Alternatively, if the general alignment of elements in the scene was accurate, a higher score was given.

#### Relative positioning

This criterion assessed how accurately elements within the scene (including scene-defining properties such as windows) were positioned relative to each other, capturing whether the arrangement of recollected elements was accurate. A low score was given if the organisation of objects recalled appeared random – i.e., suggesting that the objects were remembered without any recognition for where they appeared, meanwhile a high score was given if all elements were recollected with high accuracy in relation to each other.

#### Spatial proportions

This metric assessed the accuracy of the distances between objects and key features of the scene. If distances across recollected elements were severely distorted (i.e., across the scene objects were placed too close or too far apart, or if elements were drawn further from the image edge than was shown), a lower score was given. If the distances between objects were accurate, then a higher score was given.

#### Viewing perspective

This criterion assessed whether the overall perspective and viewpoint represented in the drawing matched the original viewing angle of the scene image. A low score indicates a significant deviation from the original viewing angle, whereas a high score suggests a near-perfect match in perspective.

#### Boundary extension

This metric aimed to capture whether participants showed evidence of drawing beyond the boundaries of the original image shown (e.g., whether the image is drawn from a more ‘zoomed out’ perspective), a previously reported distortion in scene memory paradigms (Bainbridge & Baker, 2020). A low score was given if there was evidence of boundary constriction (i.e., drawing features on a significantly larger scale), a high score indicated evidence that participants had represented the scene from an expanded viewpoint, such as drawing central features on a smaller scale. A score of 0 indicated a drawing accurate to scale.

#### False memory

This criterion assessed whether participants had recalled objects or details in their drawings that were not present in the original scene image. This may have included details drawn that were not present in the original scene image, or objects that were incorrectly duplicated (e.g., drawing an item multiple times), reflecting inaccuracies in memory reconstruction. A low score indicated the presence of no falsely recalled elements, and a high score indicated an inaccurate drawing with many falsely recalled elements.

Scores derived from this coding scheme allow us to access the level of detail provided in recollected scene drawings, including scores for object memory precision (average across three object metric scores), scores for spatial memory precision (average across two spatial metric scores), which could also be combined into an overall scores (average across all five metrics), for each participant (Fig. 1D). Coders used an online form on Qualtrics to rate each image in turn, coding one participant at a time. The order of image coding was randomised. Coders were reminded to be thorough, but fair in their judgements, and to focus on the participant’s memory of the reference scene image, rather than their artistic/drawing abilities. Each metric was scored with consideration for other scoring items, whereby if only a few items were recalled relative to the original scene image, this frequently reflected in the scores for other items. However, it was theoretically possible to score high on one metric and not the other.

All scene image drawings were coded by two coders, one of which was an author (EA), and one coder who was naïve to detailed aims of the research (ERC). All drawings were coded prior to eye tracking data being analysed. Final scores of coded variables used for analyses (overall memory, false memory, boundary extension) were averaged across both coders. The presence of correct objects recalled in scene drawings were first identified, and any discrepancies were decided by EA, following discussion with the research team. Intraclass correlation coefficients (ICC) and 95% confidence intervals were calculated in R (version 4.4; R Core Team 2024) using the ‘irr’ package to assess interrater reliability. A mean-rating (k = 2), consistency, two-way model was used, reflecting fixed raters and the goal of assessing consistency of scores across trials. Interrater reliability was good across coders for participants’ overall memory scores (0.80, 95% CI [0.75, 0.84]), object memory scores (0.77, 95% CI [0.71, 0.81]), and spatial memory scores (0.80, 95% CI [0.75, 0.84]).

### Additional measures

#### Subjective experience ratings

In addition to the scene encoding and scene drawing recall task, additional measures were included to obtain subjective ratings of participants’ experiences of the scene images during recall. All questionnaire-based data was collected using Qualtrics (https://www.qualtrics.com), and three short scale-response items were used here. Firstly, participants were asked ‘how much confidence do you have that your recalled scene drawing you produced is representative of this scene image you were initially presented?’. Responses were given on a 5-point Likert scale where 1 = ‘not at all confident’, and 5 = ‘completely confident’. Secondly, participants were asked ‘When recalling this scene image, did you visualise/remember this scene as a 2D image or as if you were immersed in the scene?’. Here, responses were given on a scale of 0-100, where 0 = ‘I had a 2D image of the scene in my mind’, and 100 = ‘I felt fully immersed in the scene, as if I was there’. Finally, participants were asked ‘How vivid was this scene in your mind when recalling it from memory? This item was rated on a scale of 0-100, where 0 = ‘I did not see anything in mind’, and 100 = ‘I had an extremely detailed image in mind’. Two additional questions pertaining to artistic abilities were also included, these referred to participant’s subjective perceptions towards their artistic ability: ‘How would you rate your artistic abilities?’ with five fixed responses, 1 – Novice, 2 – Beginner, 3 – Intermediate, 4 – Advanced, 5 – Expert, and ‘How many years of experience they have in artistic activities (e.g., art classes, college courses, hobbies)’, with fixed responses for ‘less than 1 year’, ‘1-2 years’, ‘3-5 years’, ‘6-10 years’, and ‘more than 10 years’.

### Statistical analysis

Whilst separate object and spatial scores were measured to reflect different aspects of memory recall, due to the high positive correlation between these variables (r = 0.97, see Supplementary Fig. S5), we instead calculated overall memory precision scores and did not discriminate between object and spatial memory throughout. Additionally, both total number of fixations and scanpath length were highly positively correlated (r = 0.80, see Supplementary Fig. S5), suggesting each saccade was of similar length on average. To reduce potential multicollinearity, a principal component analysis (PCA) was conducted on these two variables to create a single/combined exploration metric. Both variables were standardised, and the first principal component accounted for 88.9% of the variance and positively loaded onto fixation count and scanpath length (loading = 0.71 for both, respectively). Therefore, this component was used and inferred to represent the ‘extent of visual exploration’ during encoding *as our lower-order measure*.

#### Model 1 – Likelihood of recalling an object AOI if fixated upon at encoding

To address our first hypothesis, we fit a generalised linear mixed model (GLMM) to assess whether object AOIs fixated on during encoding were more likely to be subsequently recalled in participants’ scene drawings (Model 1). For each object AOI, binary variables indicated whether the AOI was fixated on during encoding and whether it was recalled (fixation: 0 = not fixated, 1 = fixated; recall: 0 = not recalled, 1 = recalled). The GLMM model was then fitted with AOI looked at as the fixed-effect predictor, and AOI recalled as the binary outcome. Participant and scene image were included as random effects to account for individual differences and scene specific variability. This model was fitted with a binomial distribution and logit link function, with model estimates calculated using maximum likelihood (Laplace approximation). Statistical significance was evaluated using 95% bootstrapped confidence intervals.

#### Model 2 – Relationship between ‘lower-order’ eye movement component and memory precision

To explore how the extent of eye movements during encoding may relate to subsequent scene memory performance, we fit a linear mixed-effects model (LMM) to assess whether our ‘lower-order’ visual metrics could predict memory precision scores. Here, lower-order measures reflect the intensity of exploration, such as the frequency and spatial extent of fixations. This model set to explore the relationship between the extent of visual exploration (using the PCA-derived exploration metric as a fixed-effect predictor variable) and overall memory precision scores.

#### Model 3 – Relationship between ‘higher-order’ eye movement measures and memory precision

To examine the relationship between relational eye movements and scene memory performance, we fit a final LMM to examine the relationship between percentage of refixations (out of all fixations), SGE, and GTE, and overall memory precision memory scores. These predictor variables capture the ‘higher-order’ visual metrics, which summarise how gaze is organised across time and space, reflecting more intentional spatial-based looking patterns during encoding and capturing how participants shift their gaze between regions of the scene image.

LMMs were estimated using restricted maximum likelihood (REML). Statistical significance of fixed effects was evaluated using the lmerTest package version 3.1.3 (Kuznetsova et al., 2017), which calculates p-values via Satterthwaite’s degrees of freedom approximation. Bootstrapped confidence intervals (95% level) were calculated with 100 simulations. Model fit and variance explained were assessed using marginal and conditional R^2^. Although some bootstrap samples resulted in convergence issues (i.e., singular fits), the main model did not exhibit convergence issues. Both models included random intercepts for participant and scene image to account for subject- and item-level variance. Random slopes were not fitted, as the effect of each eye tracking variable was expected to be consistent across participants and scene images. We also compared relative model fit of the lower-order (Model 2) and higher-order model (Model 3) using the Akaike information criteria (AIC), whereby a lower value indicated better model fit.

All statistical analyses were conducted in R (version 4.4; R Core Team, 2024). Both the GLMM and LMMs were fitted using the lme4 package in R (version 1.137; Bates et al., 2015). Trials whereby drawings were not recalled, or where the eye tracker failed to record sufficient samples (error > 25%) were handled as missing data in the model. The hypotheses and analyses for this study were preregistered on the Open Science Framework (OSF) and can be found here: https://osf.io/wf5nv. The current analysis included exploratory analyses that were not preregistered, including exploring whether boundary extension and false memory were predictors of subsequent memory precision scores. Additionally, two key lower-order variables (total fixation count, scanpath length) were combined into a single PCA-component to avoid potential multicollinearity (see above) following initial inspection of the data. The distinction between ’lower-order’ and ’higher-order’ eye movement measures was proposed during data collection and prior to analysis, as a theoretically motivated way of organising our pre-registered predictor variables into two classes reflecting the extent versus the organisation of visual sampling (see Introduction); this structuring was not itself specified in the pre-registration. The subsequent AIC-based comparison of the resulting lower-order and higher-order models was introduced during analysis to formally assess their relative fit and was also not pre-registered. Our memory outcome variable for Models 2 and 3 was adjusted to overall memory precision, rather than object and spatial separately due to high correlation between scores (see above). Hypotheses were therefore adjusted to account for this change. The R code for this study is available via OSF at: https://osf.io/c2xr9/overview?view_only=8e5af99b239d4e869088289f74beaa6c.

## Acknowledgements

This work was supported by a South East Network for Social Sciences (SeNSS) Economic and Social Research Council (ESRC) studentship [ES/P00072X/1; to E.A.], Royal Holloway, University of London, and the Biotechnology and Biological Sciences Research Council (BBSRC) [BB/V010549/1; to C.J.H.]. We would also like to thank James Green, for his support in the manual coding of participant drawings, and Saaria Sohail, for her involvement in the setup of this study, and the support throughout.

## Data availability

The data of this study are available via OSF at: https://osf.io/c2xr9/overview?view_only=8e5af99b239d4e869088289f74beaa6c.

## Code availability

The R and MATLAB code of this study are available via OSF at: https://osf.io/c2xr9/overview?view_only=8e5af99b239d4e869088289f74beaa6c.

