## Supplementary information for "Eye movement markers of relational encoding predict scene memory precision"

#### Supplementary Figure S1: Distributions of eye tracking variables

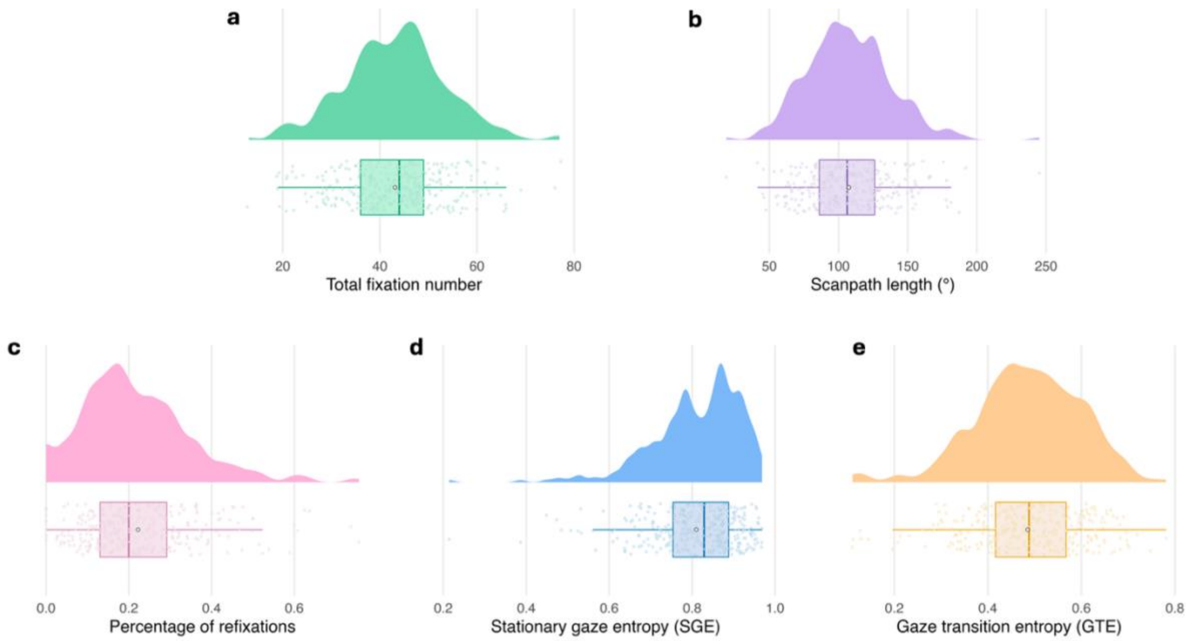


**Fig. S1. Distribution raincloud plots for lower- and higher-order eye tracking variables.** (A-B) Distribution of lower-order eye tracking variables: total number of fixations and scanpath length, respectively. (C-E) Distribution of higher-order eye tracking variables: percentage of refixations, stationary gaze entropy (SGE), gaze transition entropy (GTE), respectively.

#### Supplementary Figure S2: Distribution plots for exploratory measures

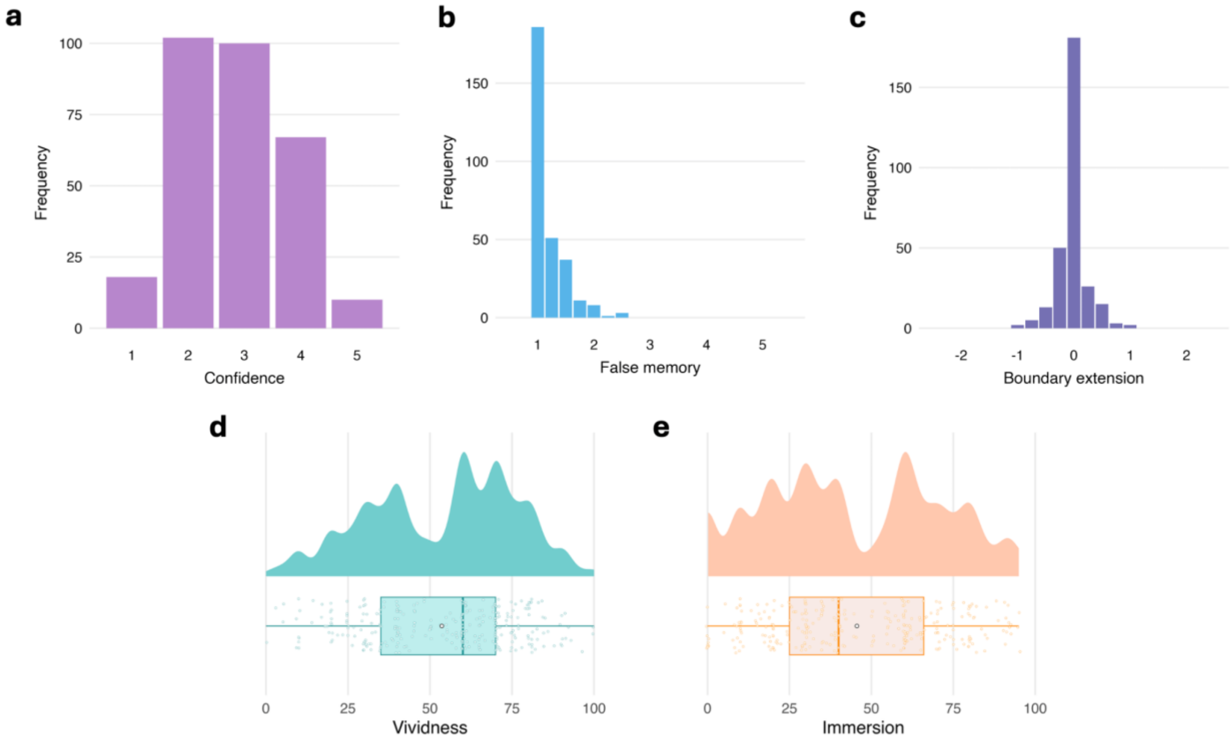


**Fig. S2.** (A-C) Distribution of discrete exploratory measures: confidence, false memory and boundary extension, respectively. Whilst confidence was the self-reported assurance one had in the accuracy of their recollected drawing, both false memory and boundary extension were obtained from manual coding of participants drawings, with little evidence of either memory error being present across participants' scene drawings. (D-E) Distribution of subjective reported feelings of vividness of the image in one's mind, sense of immersion in the image during recall.

#### Supplementary Figure S3: Linear mixed effects model (LMM) plots with subjective and exploratory predictors.

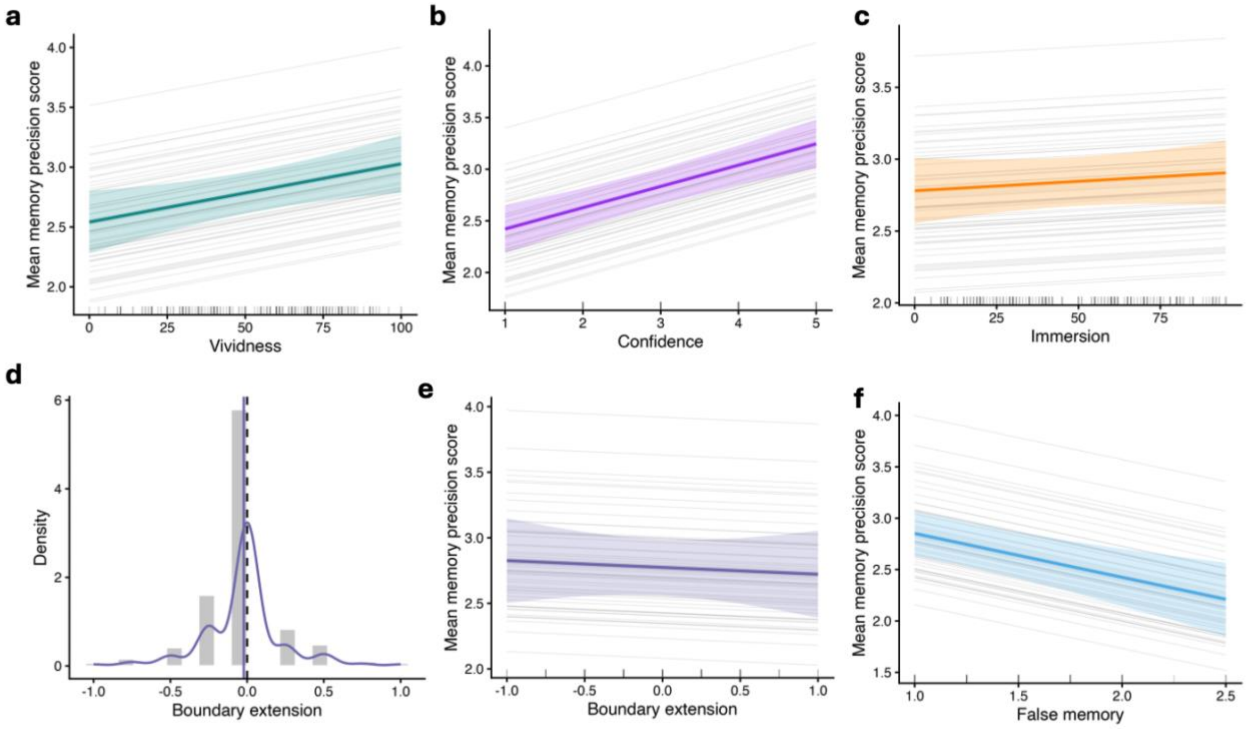


**Fig. S3. Associations between subjective memory characteristics, false memory, and boundary extension and memory performance.** (A-C). Linear mixed effects model (LMM) estimates showing the association between participants' self-reported subjective vividness, confidence, and immersion, respectively. (D) Distribution of boundary-extension scores. The dashed vertical line indicates zero boundary extension or constriction, and the solid vertical line represents the group mean. Grey bars show distribution of memory scores, and the coloured curve indicates the smoothed density. (E-F) LMM estimates showing the association between boundary extension and false memory, manually coded from participants' scene drawings. Grey lines represent individual participant relationships, coloured lines represent group-level model estimates with shaded areas showing 95% confidence intervals. Grey marks along the x-axis indicate the distribution of observed predictor values.

#### Supplementary Figure S4: Example low, medium, and high precision scene drawings with reference image and average gaze heatmaps (N = 56)

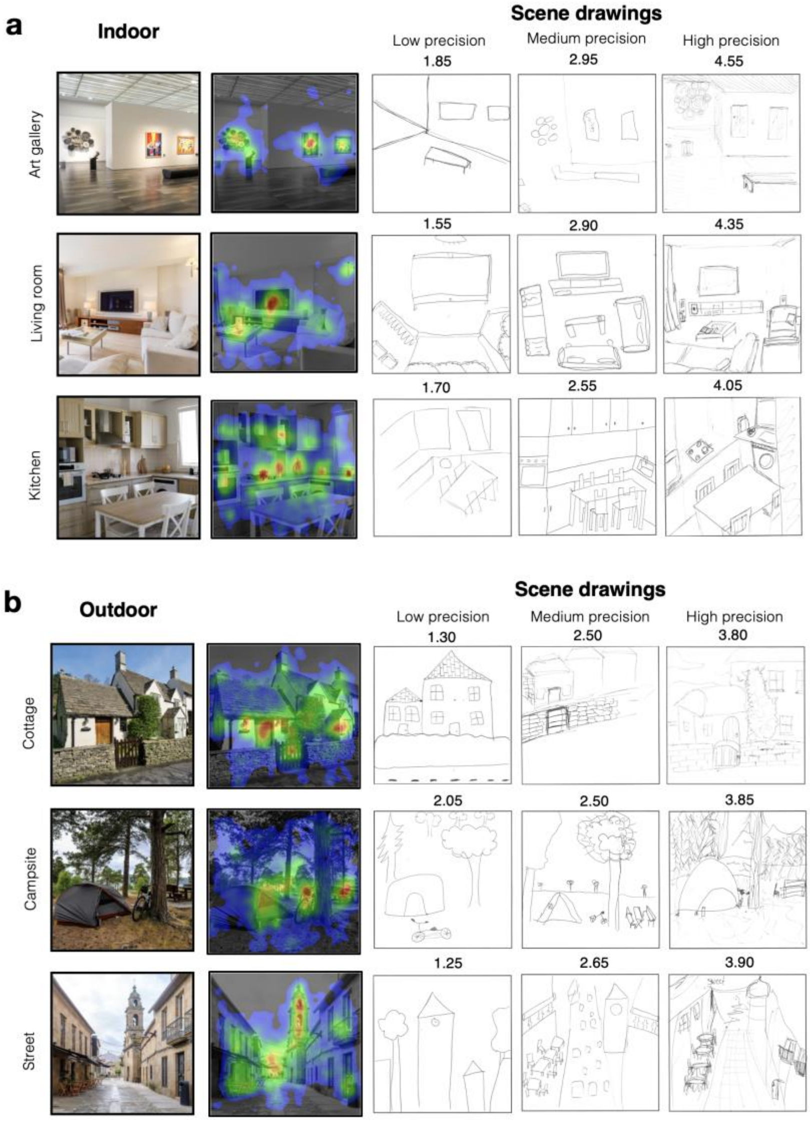


**Fig. S4.** All drawings below are from separate participants to demonstrate a variety of scene drawings from the visual free recall task for (A) indoor scene images and (B) outdoor scene images. Scene images included below are from Adobe Stock, with all images obtained under standard licensing. Drawings are presented based on 'low' 'medium', and 'high' score examples for demonstration only, and are derived from average overall memory scores (mean average of all object and spatial memory coded items). Average overall memory scores for low, medium, and high average drawings presented here range from 1.25-2.05, 2.55-2.90, and 3.80-4.55, respectively. Gaze heatmaps have red to blue gradient, representing the total number of gaze samples at each location scaled for each image, with red indicating a higher number of gaze samples.

#### Supplementary Figure S5: Correlation matrix between all key memory and eye tracking variables

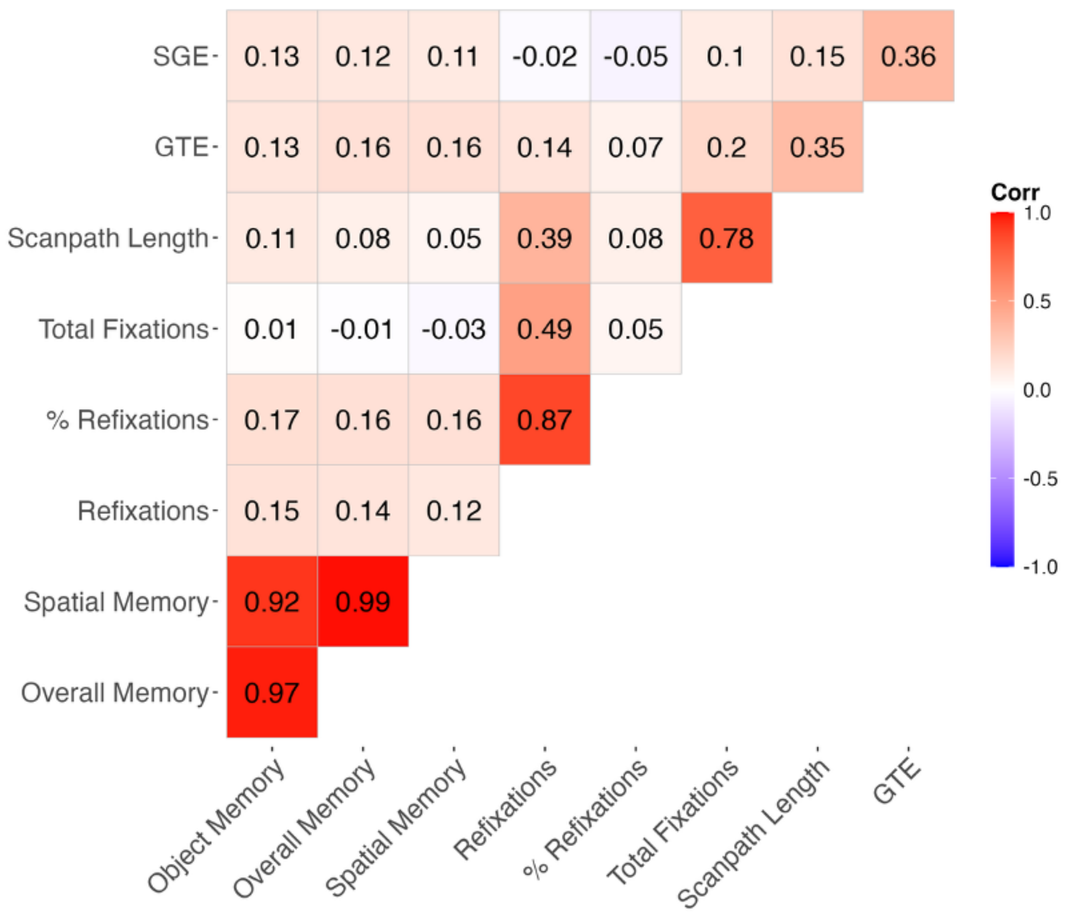


**Fig. S5. Correlation matrix between all key memory and eye tracking variables.**

### Supplementary note 1

#### Coding guide

The overall aim is to evaluate participants' memory for 6 real-world scene images. These images encompass various settings: domestic spaces such as kitchens and living rooms, and public areas such as art galleries. Each image is presented as a 500 x 500 pixel square, encompassing spatial and object details relevant to the scene category. In the task, we asked participants to draw the scene images from memory after viewing the original scene image for 20 seconds on a desktop computer. Drawing serves not only as an output of cognitive processes but also as a rich source of data for investigating cognition. The act of creating visual representation provides insight into what information was perceived, encoded, and stored in memory, as well as how this information was organised and understood. In this study, we not only want to know how much participants remember about objects, but also how well they can recall the spatial relationships between objects and the overall layout of the room. This study is rooted in the understanding that memory is not just about recalling individual items but also about how understanding and remembering how these items are related to one another in space.

As a rater, you will compare the participants' drawings to the original scene images using a sliding scale scoring system. This system has been designed to assess six main dimensions of object and/or spatial memory:

1. **Object presence and detail –** the number of objects correctly recalled from the scene image and the amount of detail remembered in those objects.
2. **Spatial distortion and rotation of objects –** the accuracy of the recalled size, shape, and orientation of individual objects.
3. **Relative positioning –** how accurately the positions of the objects and features are recalled relative to each other.
4. **Spatial proportions –** the accuracy of recalled distances between objects and features.
5. **Viewing perspective –** whether point of view of drawing matches the perspective of the reference image.

Exploratory dimensions:

1. **Boundary extension –** whether participants include detail consistent with extending image boundaries (e.g., drawing detail beyond boundary, drawing central features on significantly smaller scale, idea of whether image is further away/closer up than reference image).
2. **False memory –** the presence of objects drawn by participants that were not otherwise present in the original scene image, including duplication of objects that are not representative of the original image.

By scoring each participant's drawings across these dimensions, the study captures a more comprehensive picture of their spatial memory. This multi-dimensional approach provides a more robust and nuanced understanding of memory, reflecting that remembering involves not just recalling items but also their spatial and relational attributes.

**Instructions for Coders**
On each trial, first familiarise yourself with the original scene images so you have a solid understanding of the correct layout, object placement, and spatial proportions. You should know where each object belongs and its relative size, shape, and orientation.

**Object Presence & Detail:** When evaluating this criterion, please count the number of unique objects in the participant's drawing, and assess the level of detail included in these objects. Compare this to the total number of unique objects and object detail in the original scene image. Rate on a scale from 1 (None) to 5 (All), where a score of 1 indicates none of the original objects were included, or objects included lack recognisable detail, and a score of 5 indicates all original objects were included with substantial detail. In some cases, participants may combine objects into clusters (e.g., drawing a “study area” rather than drawing a desk, chair and filing cabinet”). These responses can be penalised but still reflects partial memory for the gist of what objects were present.

**Spatial Distortion & Rotation of Objects:** Here, evaluate the size, shape, and orientation of the individual objects within the image. If these aspects are highly distorted or misoriented compared to the original scene image, give a score of 1. If they are very accurate, assign a score of 5.

**Relative Positioning:** Here, you are evaluating how accurately objects and features (including doors and windows) are positioned relative to each other. A score of 1 should be given if none of the objects or features are placed correctly relative to others (i.e., randomly placed), while a score of 5 should be given if all objects and features are accurately positioned in relation to each other.

**Spatial Proportion:** This criterion assesses the accuracy of the distances between objects and features. A score of 1 indicates completely distorted distances, while a score of 5 suggests very accurate distances, matching the original scene drawing. For example, some individuals may draw the scene in a way object distances are compressed, or further away from the image edge than was shown in reality.

**Viewing perspective:** Here, consider whether the overall perspective and point of view depicted in the drawing aligns with the original viewing angle of the reference scene image. A score of 1 indicates a significant deviation from the original viewing angle, whereas a score of 5 suggests a near-perfect match in perspective.

Exploratory dimensions:
**Boundary extension:** Here, you are assessing whether participants have drawn beyond the boundaries of the original image shown. For example, is the overall image drawn from a more ‘zoomed out’ perspective. A score of -2 should be given if there is evidence of boundary contraction (e.g., drawing features on a significantly larger scale), a score of 0 indicates a drawing accurately to scale, and a score of 2 should be given if there is detail consistent with extended image boundaries (e.g., drawing detail beyond image boundary, drawing central features on significantly smaller scale).

**False Memory:** Lastly, this criterion assesses whether participants have recalled objects or features in their drawings that are not present in the original scene image. This may include detail drawn that is not present at all, or objects that are incorrectly duplicated (e.g., drawn multiple times that is not observed in the reference image). A score of 1 indicates the presence of no incorrect additional objects/features, and a score of 5 indicates an inaccurate drawing with many incorrect details recalled.

Object metrics: object presence, false memory
Spatial metrics: relative positioning, spatial proportions, boundary extension, viewing perspective, spatial distortion & rotation.

**Additional pointers**

- Take your time to review each participant's drawings thoroughly, and assign your scores carefully. It is important to be as fair and consistent as possible.
- If you are unsure about a score, don't hesitate to consult a fellow coder to discuss the drawing and come to a consensus.
- Record your scores for each category and for each participant's drawing using the provided sliders, and double-check your entries for accuracy before submitting them (by pressing the 'Next' button on the page).
- Remember, the goal here is not to critique artistic skill but to evaluate the accuracy of the participant's memory of the original reference scene image. Be patient, diligent, and fair in your assessments.

#### Supplementary Note 2: Self-reported artistic experience

The current study also included two forced-choice self-report questions about participants' artistic abilities and confidence in artistic skills, and their prior experience with art. These questions were collected as part of the study protocol, but were not included in the analyses presented in the manuscript. They are provided below for completeness.

**Question 1: How would you rate your artistic abilities?**
Novice: I have little to no experience or confidence in my artistic abilities.
Beginner: I have some to basic skills but lack confidence.
Intermediate: I have moderate skills and feel somewhat confident.
Advanced: I possess strong artistic skills and am quite confident.
Expert: I am highly skilled and very confident in my artistic abilities.

**Question 2: How many years of experience do you have in artisitic activities including both formal education (e.g., art classes, college courses) and informal practice (e.g., personal projects, hobbies)?**
Less than 1 year
1-2 years
3-5 years
6-10 years
More than 10 years
